# Comparative Genome Analysis of *Lactobacillus acidophilus* Isolates from Different Ecological Niches

**DOI:** 10.1101/2022.03.25.485888

**Authors:** Xudong Liu, Tongyuan Hu, Xiaoqian Lin, Hewei Liang, Wenxi Li, Xin Jin, Liang Xiao, Xiaodong Fang, Yuanqiang Zou

## Abstract

*Lactobacillus acidophilus* has been extensively applied in plentiful probiotic products and is mostly found in the gastrointestinal tract, vagina, and oral cavity of human and animal, and fermented foods. Although several studies have been performed to investigate the beneficial characteristics and genome function of *L. acidophilus*, comparative genomic analysis remains scarce. In this study, we collected 74 *L. acidophilus* genomes from our gut bacterial genome collection and the public database and conducted a comprehensive comparative genomic analysis. The analysis of average nucleotide identity (ANI), phylogenetic, gene distribution of COG and KEGG database, carbohydrates utilization, and secondary metabolites revealed the potential correlation of the genomic diversity and niches adaptation of *L. acidophilus* from different perspectives. In addition, the pan-genome of *L. acidophilus* was found to be open, with metabolism, information storage and processing genes mainly distributed in the core genome. Phage- and peptidase-associated genes were found in the genome of the specificity of animal-derived strains, which were related to adaptation of animal gut. SNP analysis showed the differences of the utilization of Vitamin B12 in cellular of *L. acidophilus* strains from animal gut and others. This work provides new insights for the genomic diversity analysis of *Lactobacillus acidophilus* and uncovers the ecological adaptation of the specific strains.

## Introduction

*Lactobacillus acidophilus* is a gram-positive, rod-shaped, plasmid-free, acidic environment preferred, prokaryotic bacteria with low GC content. Many strains of *L. acidophilus* were widely used in the processing and production of multifarious dairy products. *L. acidophilus* was generally found in the human gastrointestinal tract, oral cavity, female vagina, and fermented foods (1, 2). Several studies have demonstrated the beneficial function of *L. acidophilus* to human, such as protecting the gastrointestinal tract against the colonization of *Helicobacter pylori* (3), preventing the female vagina from the infection of *Gardnerella vaginalis* by producing antimicrobial protein (4), alleviating diarrhea (5), relieving symptoms caused by cancer (6), improving immunity (7), and lowering cholesterol (8). So *L. acidophilus* can be extensively used as a commercial probiotic product (9). Since 1972, *L. acidophilus* NCFM has been commercially used as the first probiotic, the study of *L. acidophilus* NCFM also shows that its genomic characteristics, such as two-component regulatory systems, are likely related to human health (10). But the relationship between the diversity of *L. acidophilus* with different niches was rarely explored.

The comparative genomic analysis method was widely used for investigating the genomic diversity of bacteria. Many *Lactobacillus* species, such as *Lactobacillus ruminis* (11), *Lactobacillus crispatus* (12), *Lactobacillus paracasei* (13), *Lactobacillus kefiranofaciens* (14), and *Lactobacillus helveticus* (15) have been conducted the comparative genomic analysis and the results revealed the genome features, phylogenetic relationships, differential factors associated with niche adaptation and probiotic properties vary from different strains. These studies provide clear comprehension of each *Lactobacillus* species in genetic, metabolic, and functional horizontal.

Previous studies about *L. acidophilus* were mostly focused on the probiotic functional characteristics. Comparative genomic analysis is achievable to explore the differences between the functions and beneficial mechanisms of individual strains, providing better prediction and understanding of the gene functions, genetic diversity, and the relationship with environmental adaptation. Thence, this study is dedicated to a comparative genomic analysis with comprehensive genome collection of *L. acidophilus*, excavate the genetic diversity upon similarities and dissimilarities, uncover the differential genes and molecular in-depth, and detail from the point of view of niches adaptation and the potential evolutionary mechanisms.

## MATERIALS AND METHODS

### Genome collection of 74 *Lactobacillus acidophilus* strains

The cultivation, whole-genome sequencing, and de novo assembly of 17 *L. acidophilus* strains were according to our previous work for constructing a human gut bacteria collection (16). 57 *L. acidophilus* genomes were downloaded from the National Centre for Biotechnology Information (NCBI) (https://www.ncbi.nlm.nih.gov/). In total, 74 *L. acidophilus* genomes were included in this study (Table 1).

All 74 genomes of *L. acidophilus* were renumerated according to their isolate source with the first few letters of the genomes of all strains, “A” represents Animal origin; “C” represents Commercial origin; “F” represents Food origin; “H” represents Human origin, but not sure if it is of human gut origin; “Hi” represents Human infant gut origin; “Hg” represents Human gut origin; “HgS” represents Human gut origin from our collection in this work.

### The Average Nucleotide Identity (ANI) Analysis

FastANI (v1.32) (17) was used to calculate the paired ANI values matrix, and R function ‘pheatmap’ (18) was used to conduct the clustering and visualization of ANI values.

### Phylogenetic Analyses

The phylogenetic tree was inferred using GTDB-Tk (The Genome Taxonomy Database Toolkit, v1.7.0) (19) for the clustered genomes, with *Alicyclobacillus acidiphilus* NBRC 100859 (taxoid:1255277) used as an outgroup. The phylogenetic tree was visualized in iTOL (http://itol.embl.de/) (accessed on 10 August 2021) online.

### Functional annotation and analysis

Gene prediction and Clusters of Orthologous Groups (COG) classification were performed with Prokka (v1.14.5) (20), and further COG annotation was performed with the COG database. KofamKOALA (v 1.3.0) (21) was used to acquire Kyoto Encyclopedia of Genes and Genomes (KEGG) identities, and further multi-level information was annotated with the KEGG database. COG and KEGG orthologs were counted in multi-level categories. The number of gene copies at the functional and molecular levels in the COG and KEGG database was calculated using the in-house developed Python/R scripts. The heatmaps were created by Pheatmap (18). The method of ANOVA (Analysis of Variance) was two-factor without replication ANOVA (22).

### Pan-genome analysis

Pan-genome analysis was conducted using PEPPA (23), curve fitting of pan-genome was used Heaps’ law model on rarefaction and permutation (24). In-house developed Python/R scripts and micropan (25), ggplot2 (26), tidyr (27) R-package were used to conduct fitting, and accumulation curve visualizes. Unique genes of animal-derived strains were found in the present/absent matrix by PEPPAN.

### Analysis of Carbohydrate-Active enzymes

The carbohydrate metabolic enzymes of 74 *L. acidophilus* genomes were annotated by dbCAN2 (28), and the carbohydrate-active enzymes were analyzed by Carbohydrate Active Enzymes Database7 (CAZy, accessed on 10 February 2021) (http://www.cazy.org/) (29). Only the CAZY families predicted by two of dbCAN2’s three algorithms (HMMER Hotpep and DIAMOND) were included in the following analysis. The heatmaps were created by Pheatmap (18).

### Prediction of Secondary Metabolites Biosynthesis Gene Clusters

The Biosynthesis Gene Clusters were predicted by using anti-SMASH (30), phylogenetic tree was analysis by the secondary metabolites biosynthesis gene sequence similarity. The functions of secondary metabolites were interpreted according to the previous studies (31-34).

### Analysis of Single Nucleotide Polymorphism

The Single Nucleotide Polymorphism (SNPs) of *L. acidophilus* was called by Parsnp (35) against *L. acidophilus* NCFM with a complete genome used as the reference. SNPs were annotated by SnpEff (36) with “-only CDS” options, only SNPs located at CDS (Coding Sequence) were analyzed. The protein’s three-dimensional structure was predicted by SWISS-MODEL (37).

## RESULTS

### Information of Source and Genome Characteristics of *L. acidophilus*

In this study, we selected 17 genomes of *L. acidophilus* in our bacterial collection, which were isolated from the human gut in Shenzhen, Guangdong, China. The other 57 were publicly available *L. acidophilus* genomes from NCBI. Among these genomes, 2 genomes were from animal gut, 16 from commercial products, 15 from food, 30 from the adult human gut, 6 from infant gut, and 7 genomes were missing the source information. The genome size varied from 1.85 Mb to 2.09 Mb with an average size of 1.97 Mb. The GC content ranged from 34.2% to 35.7% with an average GC content of 34.6%, only strain P2 was exceeded 35%. The gene number rise evenly from 1,856 to 2,084. The average CDS number is 1,914 (Table 1) 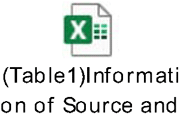.

### Genome comparison of *L. acidophilus* based on ANI

ANI analysis was a genome-based relatedness method for assessing genetic distance (38, 39), which 95% of ANI value being suggested as the delimitation criterion for species (17, 40). The result showed that most of the ANI values of 74 genomes of *L. acidophilus* were higher than 99.9% (Supplementary Figure 1), indicating these genomes shared highly similar genome sequences. Among these, strain YT1, derived from animals, share a relative farther genetic distance with other strains, which the ANI values were below 99.2%. The relationships between ANI with the source of *L. acidophilus* were shown in Supplementary Figure 1, we found strains “HgS” and strains “C” tend to cluster together, suggesting that commercial strains may also have originated in the human gut. In contrast, *L. acidophilus* strains from other sources were more scattered.

### Phylogenetic Analyses of *L. acidophilus*

To analyze the phylogenetic relationship of 74 *L. acidophilus* strains, the phylogenetic tree was constructed on the whole genomes of 74 *L. acidophilus*, which was rooted with *Alicyclobacillus acidiphilus* NBRC 100859 as an outgroup (Fig. 1). Due to the high similarities of ANI values between 74 *L. acidophilus* strains, it is difficult to determine the evolutionary relationship of *L. acidophilus*. As shown in Fig. 1, the phylogenetic tree mainly formed 4 subbranch, but genomes from varied sources were distributed in different subbranch. The biggest subbranch included the strains mainly from“Commercial”, “Food”, “Human gut”, and “Human infant”. This could indicate no significant correlation between the phylogenic relationship of strains with the different source.

**Fig. 1.**
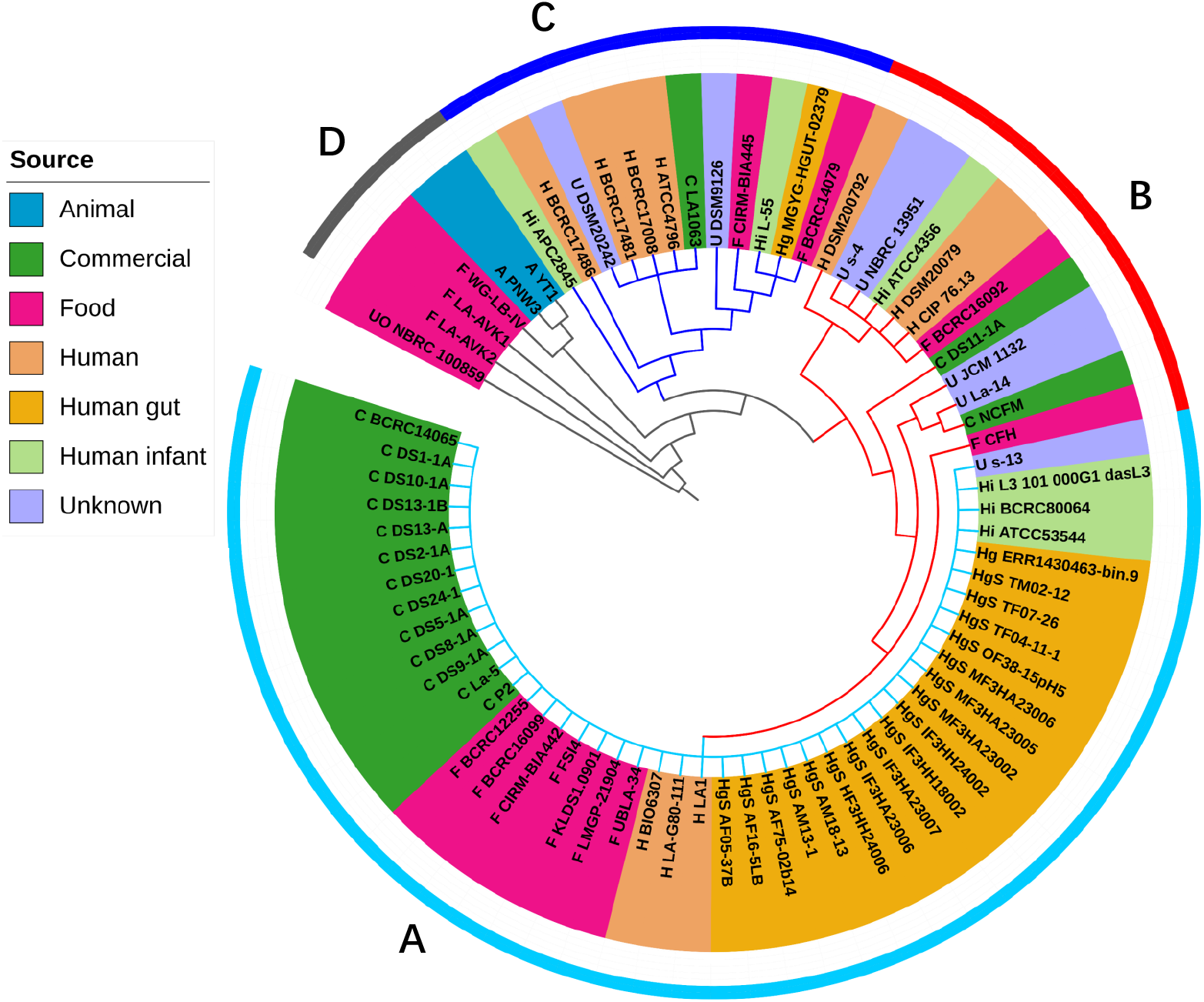
Phylogenetic analysis of *Lactobacillus acidophilus*, Phylogenetic tree was constructed based on whole-genome, imply the putative correlation between the diversity and source of 74 strains of *Lactobacillus acidophilus* with *Alicyclobacillus acidiphilus* NBRC 100859 as an outgroup. The resulting phylogenetic tree divided those strains in four clades (clades A to D). Clade A consisted of half of strains from human, food, and commercial source; clade B gathered 12 strains, among which two strains from food, two strains from commercial products, four strains from human, four strains from unknown source; clade C included 12 strains, which were 7 strains from human, one strain from commercial products, two strains from food, two strains from unknown source; while clade D consisted of five strains, among which two isolates were from animals, three isolates were from unknown source.

### Genomic Function Analysis of *L. acidophilus*

The distribution of functional classification of genes based on the COG and KEGG annotation of 74 *L. acidophilus* strains were showed in Supplementary Figure 2 and Supplementary Figure 4. The genes are mainly distributed in Carbohydrate transport and metabolism (G), Translation, ribosomal structure and biogenesis (J), and Amino acid transport and metabolism (E) for COG categories (Supplementary Table 1. 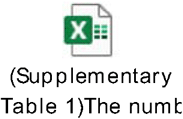 According to the KEGG annotation, the genes are mainly distributed in genetic information processing, signaling and cellular processes, and carbohydrate metabolism (Supplementary Table 2) 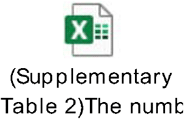, that were consistent with the categories of COG. In addition, vitamin B12 import ATP-binding protein BtuD possess the most numbers of the genes in the COG categories (Supplementary Fig 3), which was a part of the ABC transporter complex BtuCDF involved in vitamin B12 import and energy coupling to the transport system. Nevertheless, the molecule with the most gene numbers in the KEGG database is sugar PTS system EIIA component (Supplementary Fig 5), a phosphoenolpyruvate-dependent sugar phosphotransferase system (sugar PTS), involved in catalyzing the phosphorylation of incoming sugar substrates concomitantly with their translocation across the cell membrane (41). We next analysis the different function categories in 74 *L. acidophilus* strains according the COG and KEGG annotations. The top 30 different molecules in COG categories were consist of many tRNA and rRNA related genes. Interestingly, Vitamin B12 import ATP-binding protein BtuD was found in the top 30 different molecules (Supplementary Fig 3). The top 30 distinct molecules in KEGG categories contained the kinases, epimerases, transposases, mutases, transport proteins, etc. The most different molecule was wecB: UDP-N-acetylglucosamine 2-epimerase (Non-hydrolyzing) (Supplementary Fig 5). Additionally, no obvious correlation between the different function categories of strains with the source.

### Pan-genome analysis of *L. acidophilus*

The pan-genome analysis is performed for investigating the distribution of all the genes at the species level. In this work, the boxplot, fitting curve, and accumulation curve were generated from the pangenome data and shown in Figure 2. The boxplot and fitting curve reflected the number of genes increases with the addition of new genomes. The pan-genome of *L. acidophilus* remained unsaturation with the Heap’s alpha being below 1.0 (Fig. 2 A). According to the previous study (23), strict core genes are found in all genomes, core genes are found in 99%-100% of genomes, soft core genes are found in 95–99% of genomes, shell genes are found in 15–95%, while cloud genes are presented in less than 15% of genomes. The obtained data showed that there were 1,573 strict core genes, 0 core genes, 221 soft core genes, 92 shell genes, and 443 cloud genes (Supplementary Table 3) 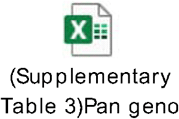.

**Fig. 2.**
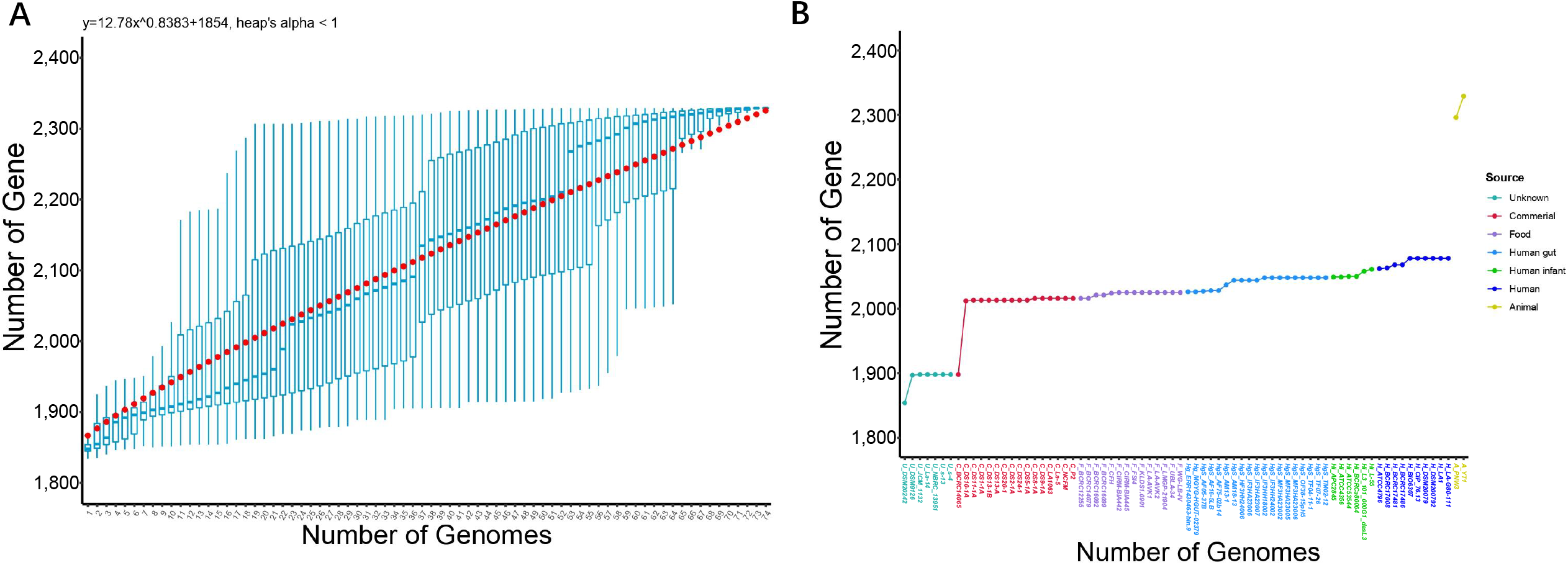
**A**, The pan-genome of *Lactobacillus acidophilus*. Boxplot is represented by the accumulated number of new genes against the number of genomes added. The fitting curve is fitted according to the median value of the permutation. **B**, The pan-genome accumulation of *Lactobacillus acidophilus*. The increase in the pan-genome accumulation curve is on the basis of the gene of the previous genome.

The outcomes of the distribution of the functional categories based on KEGG and COG annotation of core genome, accessory genome, and unique genome were depicted in Supplementary Fig 6 and Supplementary Fig 7, respectively. Metabolism, information storage, and processing genes were mainly categories. Moreover, the core genome encodes more conservative genes involved in translation, ribosomal structure and biogenesis than [J], amino acid transport and metabolism [E], and intracellular trafficking, secretion and vesicular transport [U] compared with the accessory genome and unique genome (Supplementary Fig 6). On the contrast, defense mechanisms [E], replication, recombination and repair [L], carbohydrate transport and metabolism [G], nucleotide transport and metabolism [F], and general function prediction only [R] were more abundant in the accessory genome and unique genome. For the distribution of KEGG functional categories, metabolism was the most abundant categories both in core genome, accessory genome, and unique genome (Supplementary Fig 7). The distribution of subcategories of KEGG were consistent with the results of COG.

The genome accumulation curve showed that less genes gain with the newly increased genomes from the most of different source, but the two animal-derived *L. acidophilus* strains contributed over 200 new genes (Fig. 2 B), suggesting that the animal-derived strains may encode more unique genes involved in the adaptation of the animal intestinal environment. Then we annotated these unique genes of the animal-derived strains by the COG database and found that most of these were distributed in [K] Transcription and [L] Replication, recombination and repair category for information storage and processing (Fig. 3 A). These unique genes were belonging to phage-associated, peptidase-associated, ABC transporter-associated, transposase-associated, CRISPR-associated, glycosyl transferases, transcriptional regulator, and glycosyl hydrolase related genes (Fig. 3 B), that were considered to be involved in the adaptation and evolution to the complex animal intestinal environment of *L. acidophilus*.

**Fig. 3.**
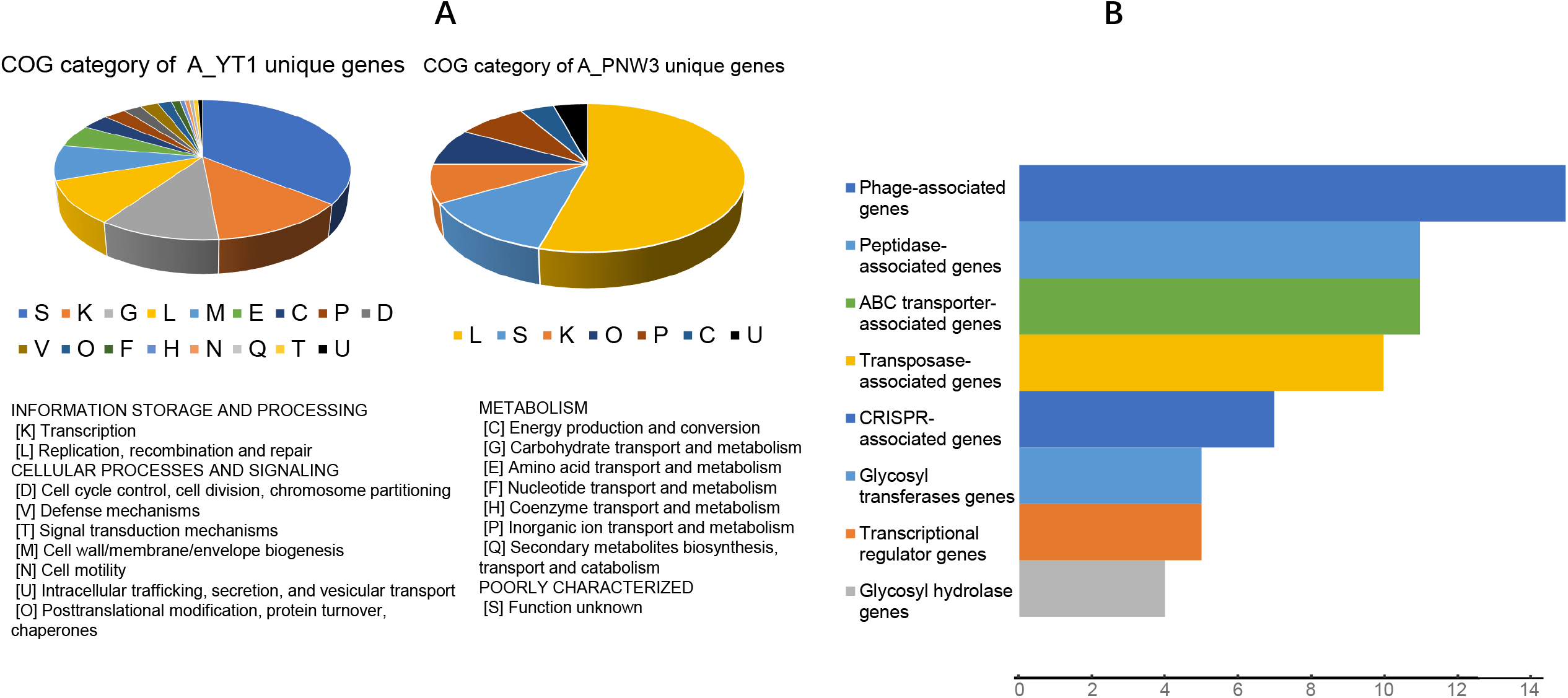
**A**, COG category of the unique genes from animal-derived strains. **B**, The recognizable and classified unique genes of animal-derived strains. Clustered bar plots showing the classification of unique genes of animal-derived strains.

### Comparative Analysis of Carbohydrate-Active Enzyme

30 CAZY families were predicted in the genome of *L. acidophilus*, which include 14 GH (glycoside hydrolases), 10 GT (glycosyl transferases), 3 CBM (carbohydrate-binding modules) families, and 3 CE (carbohydrate esterases). GH and GT are the largest number of enzymes in the carbohydrate metabolism of *L. acidophilus*. In all CAZY families, GH1 (β-glucosidase families), GH13 (α-amylase families), and GT2 (cellulose synthase families) were the most abundant CAZY families. In contrast, GH78 (α-L-rhamnosidase families), CE1 (acetyl xylan esterase families) and CE7 (acetyl xylan esterase families) were the low abundance CAZY families. The other CAZY families are distributed similarly in different strains (Supplementary Fig 8). Altogether, no significant differences were found in the genome of these 74 *L. acidophilus* strains, suggesting the carbohydrate metabolism were evolutionary conserved, even in different environmental niches.

### Prediction of the Secondary Metabolites Biosynthetic Gene Clusters in *L. acidophilus*

Bacterial secondary metabolites are alternative sources of novel drug leads. The secondary metabolite biosynthetic gene clusters (SMBGs) predicted in the bacterial genome will provide crucial information for the efficient discovery of novel natural products (42). We used anti-SMASH (30) to predict the secondary metabolite from the genomes of 74 *L. acidophilus* strains. Gassericin T, a hydrophobic class II bacteriocin with high heat stability (121°C, 10 min), good pH tolerance (pH 2–11), was strong bactericidal against many gram-positive bacteria, and thus are expected to be effective food preservatives (31, 32). We found Gassericin T was predicted in most of *L. acidophilus* genomes (73/74), and the structure of Gassericin T gene clusters of 73 strains are quite similar and no significant phylogenic relationship between the Gassericin T from *L. acidophilus* with the different source (Fig.4), indicating all these 73 strains can be used as an alternative producer for Gassericin T. Class-I lanthipeptide, a ribosomal synthesized and post-translationally modified polycyclic peptide, was considered a novel potent antimicrobial compound (33, 34). Interestingly, we discovered the Class-I lanthipeptide was only predicted in strain A_YT1 (animal origin), suggesting the animal derived *L. acidophilus* may display different antibacterial characteristics compared with the *L. acidophilus* from other niches.

**Fig. 4.**
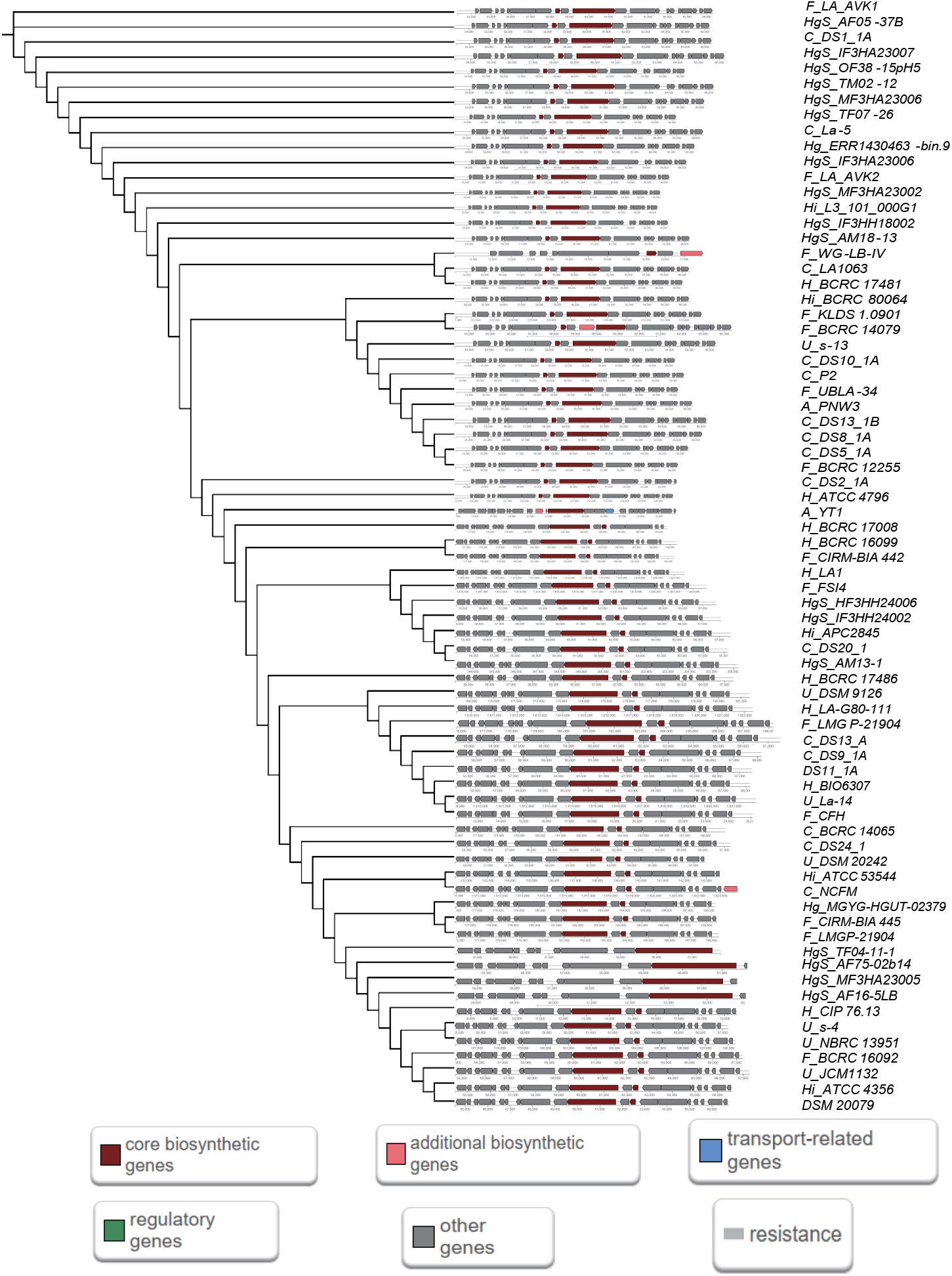
The phylogenetic tree and structure of secondary metabolite biosynthetic gene clusters of 74 strains. The secondary metabolite biosynthetic gene types were annotated by different color.

### cPotential Genetic Factor Related to Adaptation of Various Niches by Single Nucleotide Polymorphism (SNP) Analysis

SNP analysis enables descry differences among different genomes at the single-nucleotide level and uncover the genetic factor in adapting different niches of *L. acidophilus*. In total, 17,001 SNPs were identified with the number varied from 73 to 7,796, the SNPs number of the strains from the same source is close (Fig. 5 A), and most of the SNPs (7,796) were found in strain A_YT1 (Supplementary Table 4) 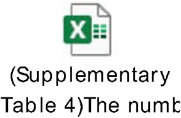, which the number was significantly more than other strains, indicating that more variations were occurred for the strain A_YT1 to adapt to the animal gastrointestinal tract. We subsequently analysis the involved genes of SNPs and found the most numerous SNPs were located in the btuD, a gene encoding Vitamin B12 import ATP-binding protein BtuD, a part of the ABC transporter complex BtuCDF involved in vitamin B12 import, responsible for energy coupling to the transport system (Fig. 5 B). Coincidentally, according to the result of COG classification at the molecular level, Vitamin B12 import ATP-binding protein BtuD was the most numbers of the genes in the COG categories in *L. acidophilus*. We furtherly investigated the highest impact of SNP variant of the btuD gene and found this SNP is a nonsense variant from the animal-derived strain YT1, which the variant was possibly pertinent to the ecological niche of this *L. acidophilus* strains. A transversion at the 918^th^ nucleotide from A (adenine) to C (cytosine) in the non-coding strand was found in the btuD gene, generating the transversion at the corresponding position from T (thymine) to G (Guanine) in the coding strand. Consequently, this transversion leads to a translation termination at the 306^th^ amino acid, tyrosine. Structurally, Vitamin B12 import ATP-binding protein BtuD was composed of NBD (nucleotide-binding domain) and TMD (transmembrane binding domain). NBD for binding and hydrolyzing ATP, provides energy for importing of Vitamin B12. TMD is primarily involved in the identification and transit of substrates through the lipid bilayer (43). Subsequently, we predicted the protein structure including the pre-mutation and post-mutation DNA sequences and found that the premature stop gained can induce the loss of NBD, thereby mediating the inactivation of the transport function of the Vitamin B12 import ATP-binding protein BtuD to stop the intracellular transport and utilizing of vitamin B12 (Fig.5 C). We hypothesized this situation may happen in the exposure of low Vitamin B12 level and *L. acidophilus* will reduce the expression and utilization of Vitamin B12 import ATP-binding protein BtuD to save the raw material for the biosynthesis of BtuD.

**Fig. 5.**
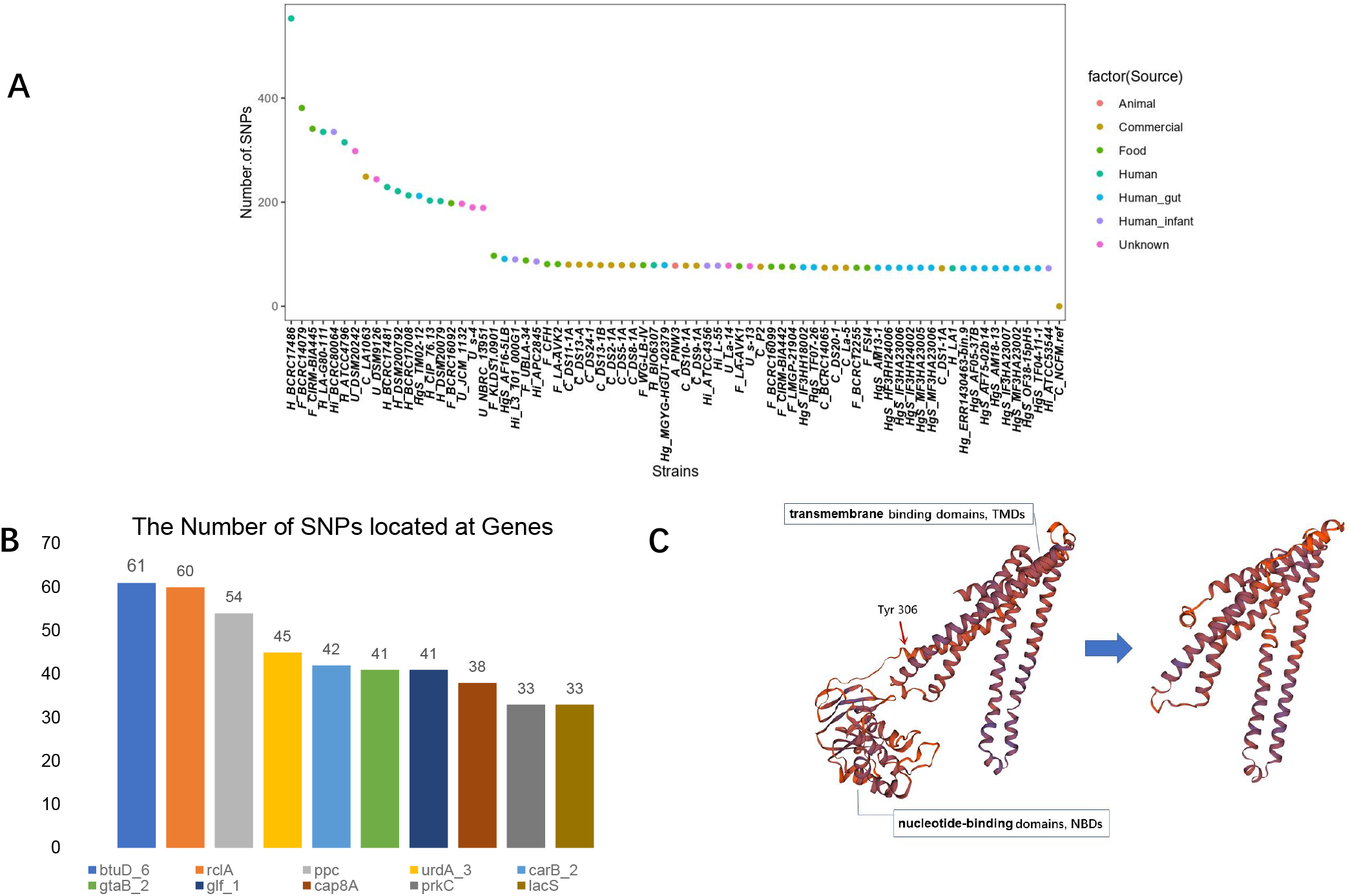
**A**, The number of SNPs in 73 strains of Lactobacillus acidophilus except A_YT1(cause too many SNPs found in A_YT1). **B**, The recognizable top 10 gene that most SNPs located. **C**, The three-dimensional structure of Vitamin B12 import ATP-binding protein BtuD, showing the difference between BtuD in normal and its mutant.

## DISCUSSION

*L. acidophilus* is a common species of lactic acid bacteria, widely used in probiotic products for many years because of its exceptional health-related benefits. The genome G+C content of *L. acidophilus* was below 36%, which can be considered as low GC bacteria (Supplementary Table 5) 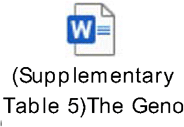. In the previous research, the probiotic properties of *L. acidophilus* have been well recognized (10). Nonetheless, the features of *L. acidophilus* to niches adaptation and evolutionary mechanism remains unexplored. In this study, we firstly reported the potential relationship among the dissimilarity of *L. acidophilus* strains with adaptation of different niches from an evolutionary perspective systemically. The genomic and metabolic characteristics of bacterial strains within species from the same source tend to be close and similar but cause differences when adapting to different environments.

By compared genome analysis of *L. acidophilus*, we demonstrate that there was no closely correlation between the diversity of *L. acidophilus* with different niches according the phylogenetic and function analysis. Interestingly, ANI clustering showed animal-derived strains shared a relative lower value with other strains, which the distinctive features were also reflected in the biosynthesis of SMBGs, pan genome analysis. We found the animal-derived strains always seem to contribute more unique genes for the pan-genomes (Supplementary Table 5). Hence, the open pan-genome denoted the genetic diversity, and the sample capacity of *L. acidophilus* could be further enriched.

What is worth mentioning is that the btuD gene is not only the most numerous SNPs located but also the most numerous gene that was predicted by the COG database at the molecular level. Therefore, we postulate that *L. acidophilus* expresses several functional Vitamin B12 import ATP-binding protein BtuD in the cell membrane to transport Vitamin B12 intracellular when the Vitamin B12 is abundant in the living environment, and it will incapacitate Vitamin B12 import ATP-binding protein BtuD by the nonsense SNP variant when the Vitamin B12 is scarce. This nonsense SNP variant was found in animal-derived strain YT1 isolated from the rat intestinal tract, which may imply the strain YT1 was original resident the rat gut under the pressure of deficient in Vitamin B12.

This is the first systematically study of *L. acidophilus* at the gene and single-base level, and revealed the characteristic of niches adaptation of *L. acidophilus*, particularly embodied in the excavation of unique genes in the pan-genome of animal-derived strains and the hypothesis on Vitamin B12 by SNP analysis. Despite this difference, the researches of this study were executed in silico. Further laboratory experiments should be conducted to explore it in transcriptome and protein translation level.

## CONCLUSION

From the results of ANI values, phylogenetic analysis, gene distribution of COG/KEGG database, carbohydrates utilization, and secondary metabolites. 74 *L. acidophilus* strains are primarily similar in general. Predictably, there are potential correlated trajectories between diversifications with niches adaptation in some ways. Besides, *L. acidophilus* was pan-genome open and more unique gene were found in animal-derived strains, one of its possible evolutionary mechanisms of niches adaptation was speculated to be associated with Vitamin B12. Thereby, the comparative genomic analysis enabled this study to exploit and better understand the biological diversity of *L. acidophilus*.

## AUTHOR CONTRIBUTIONS

Conceived and designed the experiments: Y.Z. Performed the experiments: X.L. Analyzed the data: Y.Z., X.L, L.X., T.H., W.L. and H.L. Contributed reagents/materials/analysis tools: Y.Z., X.J., X.F., and L.X. Wrote the paper: X.L. Revised the paper: Y.Z. All authors commented on the manuscript.

## DATA AVAILABILITY STATEMENT

The data that support the findings of this study have also been deposited into CNGB Sequence Archive (CNSA) of China National GeneBank DataBase (CNGBdb) with accession number CNPhis0003415.

## ACKNOWLEDGEMENTS

This work was supported by grants from National Natural Science Foundation of China (No. 32100009) and Natural Science Foundation of Guangdong Province, China (No. 2019B020230001). We also thank the colleagues at BGI-Shenzhen for sample collection, and discussions, and China National GeneBank (CNGB) Shenzhen for DNA extraction, library construction, and sequencing.

## Supplementary Material

**Supplementary Fig 1.**
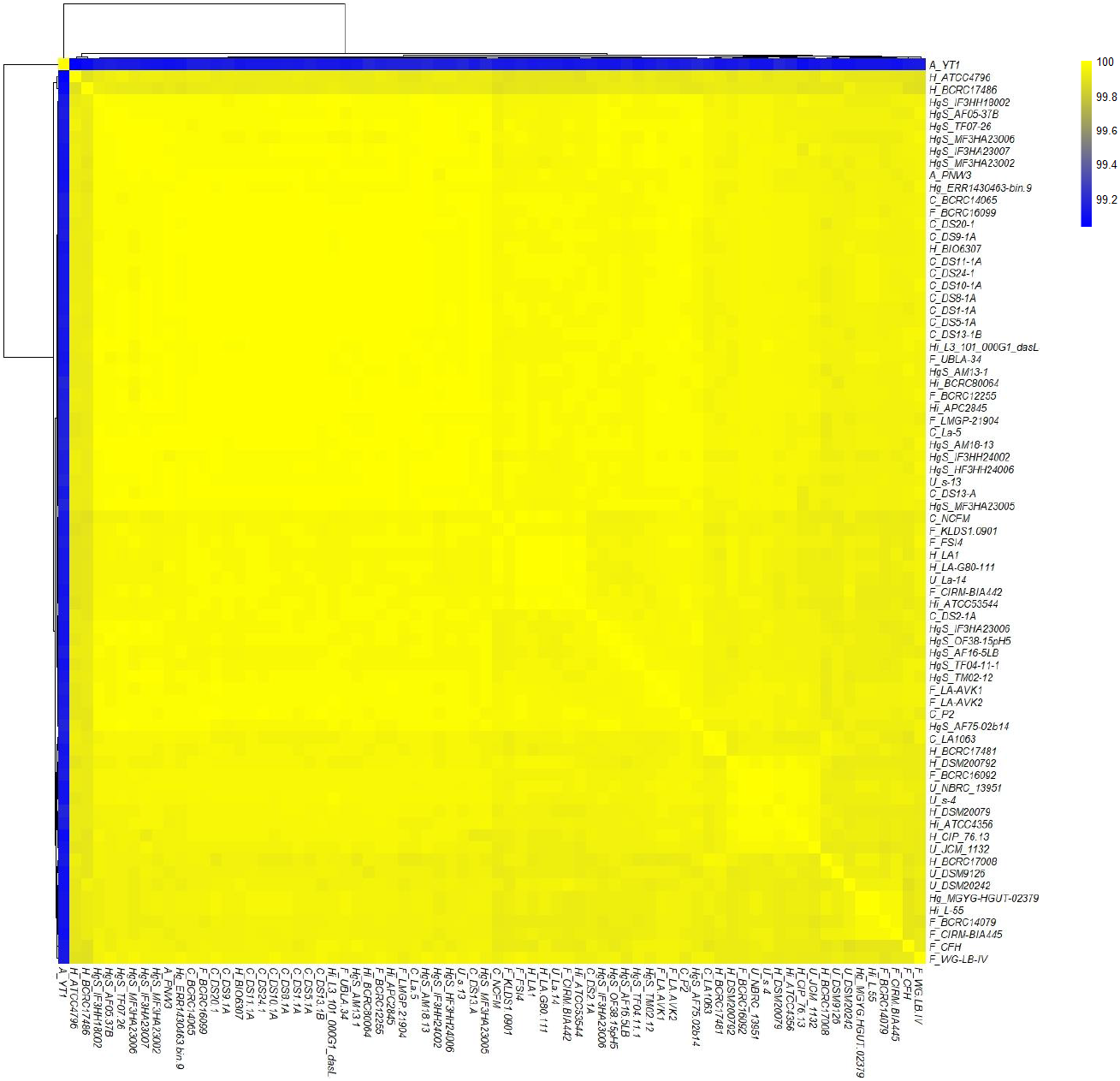
ANI heatmap of 74 strains of *Lactobacillus acidophilus*, more than 95% of ANI values was suggested as the delimitation criterion for same species, showing identity within 74 strains of *Lactobacillus acidophilus*.

**Supplementary Fig 2.**
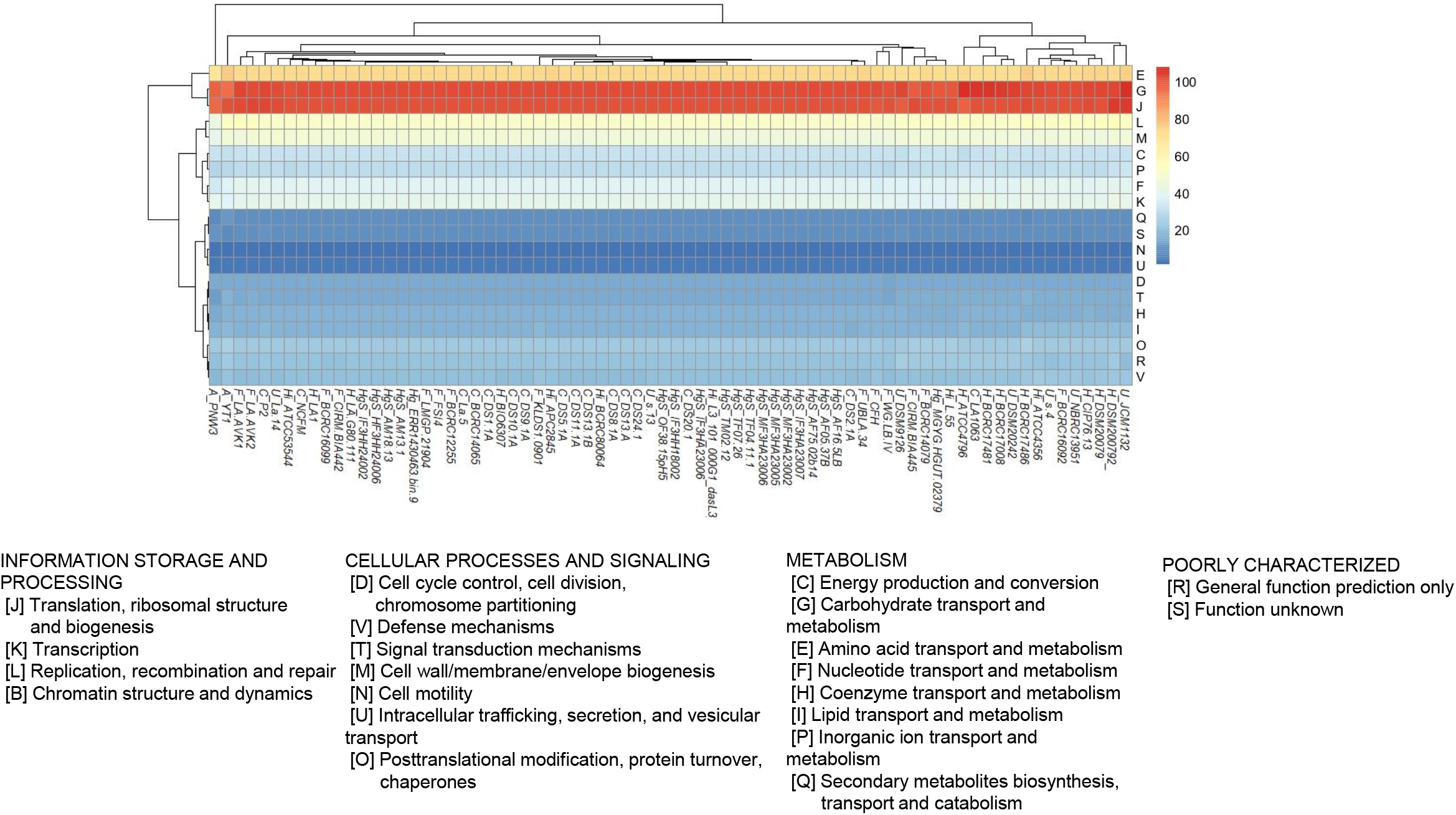
The number of genes of each strain of *Lactobacillus acidophilus* that predicted by using the COG database at the functional level. Heatmap shows the genes’ distribution at the functional level of the COG database.

**Supplementary Figure 3.**
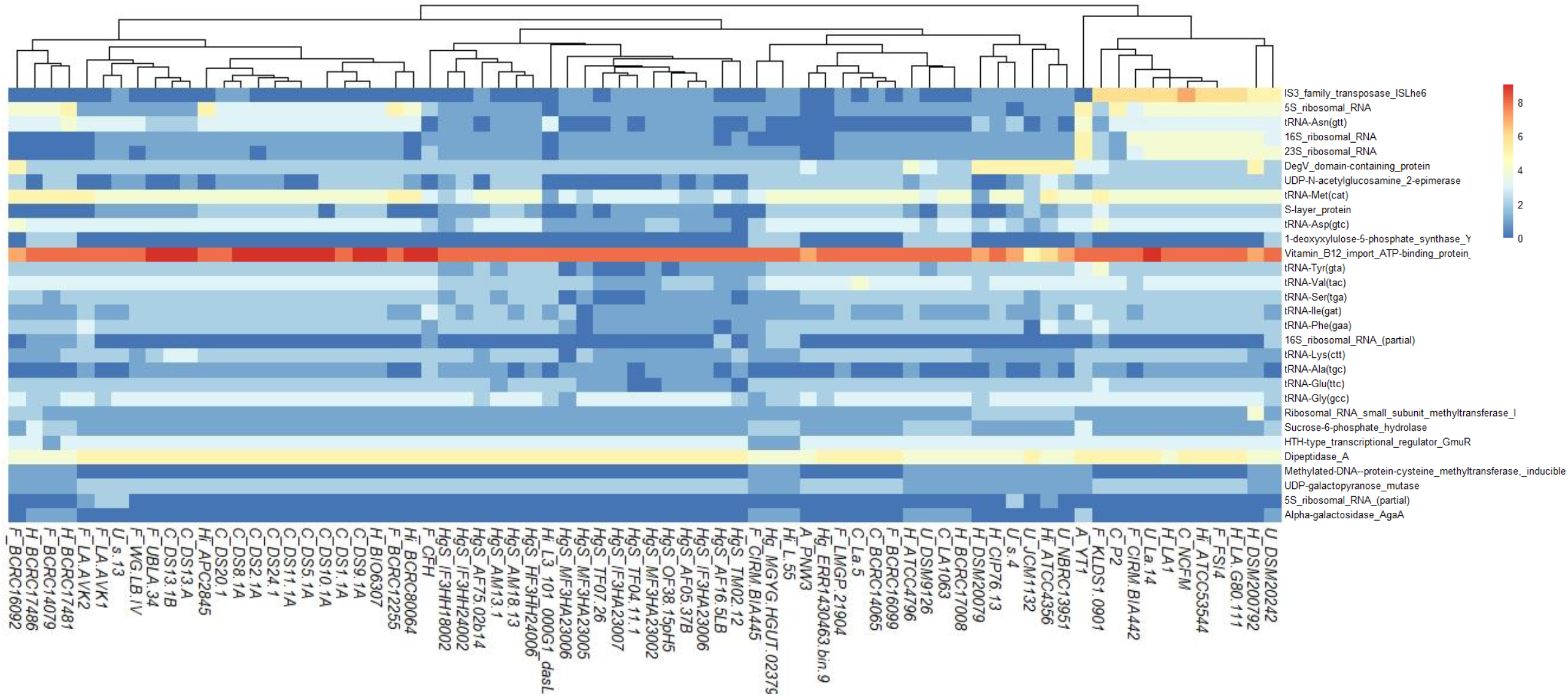
The number of top 30 variant genes annotated by using the COG database at the molecular level. The variances of COG molecules (row names) decrease from top to bottom.

**Supplementary Figure 4.**
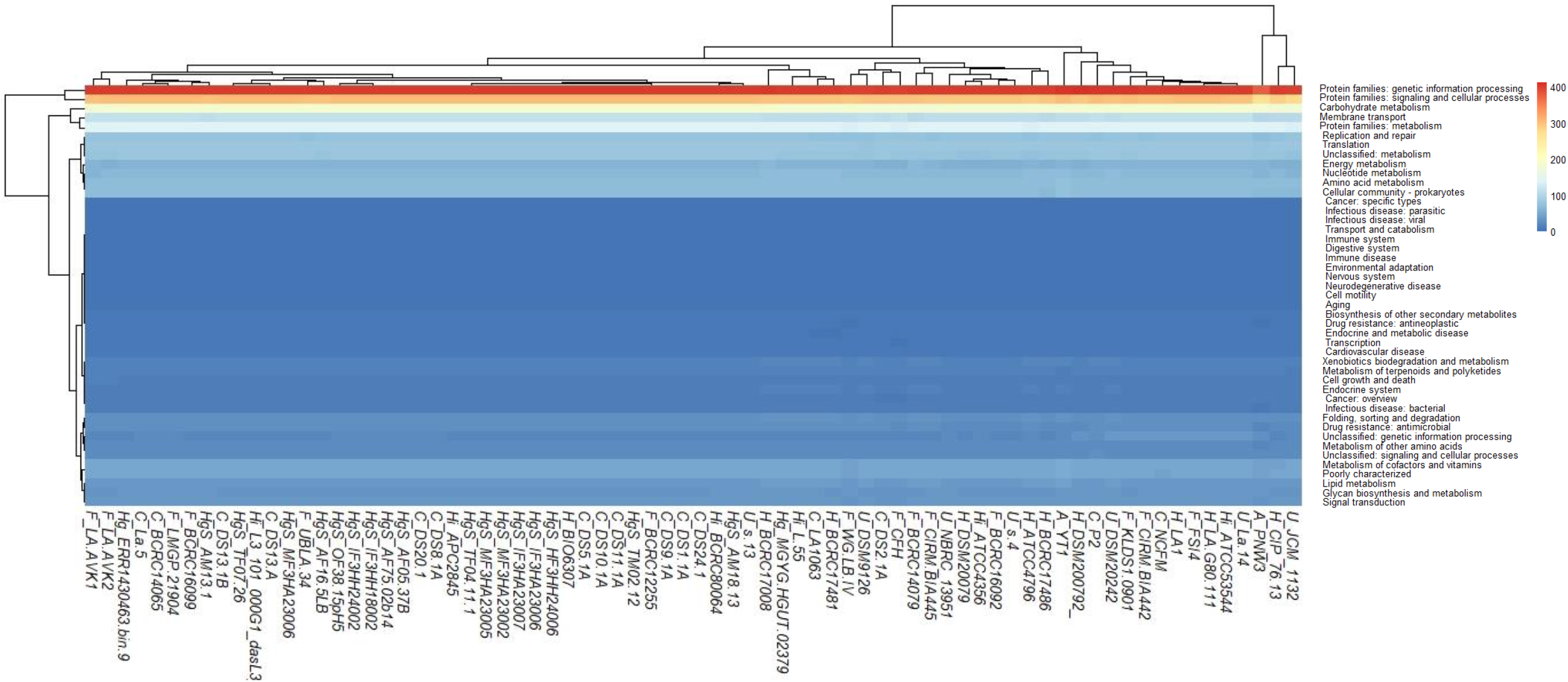
The number of genes of each strain of *Lactobacillus acidophilus* that predicted by using the KEGG database at the functional level. Heatmap showing the distribution of KEGG at the functional level.

**Supplementary Figure 5.**
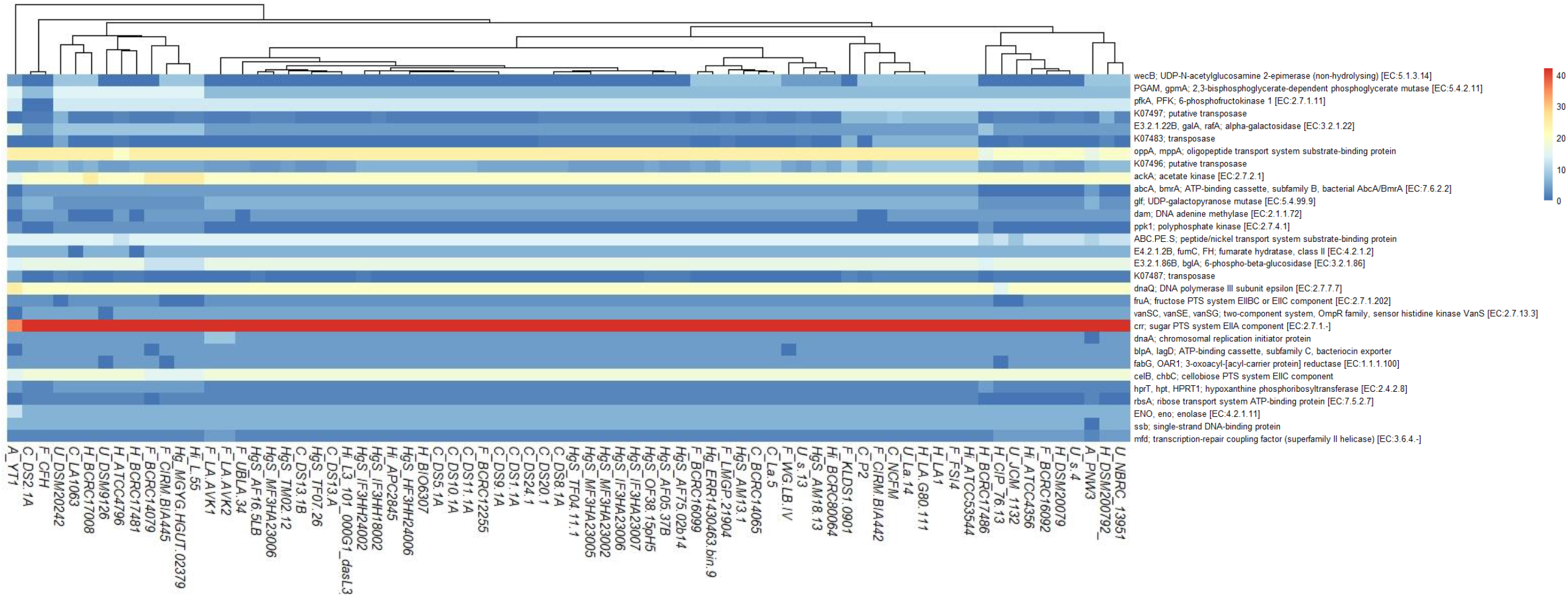
The number of top 30 variant genes annotated by using the KEGG database at the molecular level. The variances of the KEGG molecules (row names) decrease from top to bottom.

**Supplementary Figure 6.**
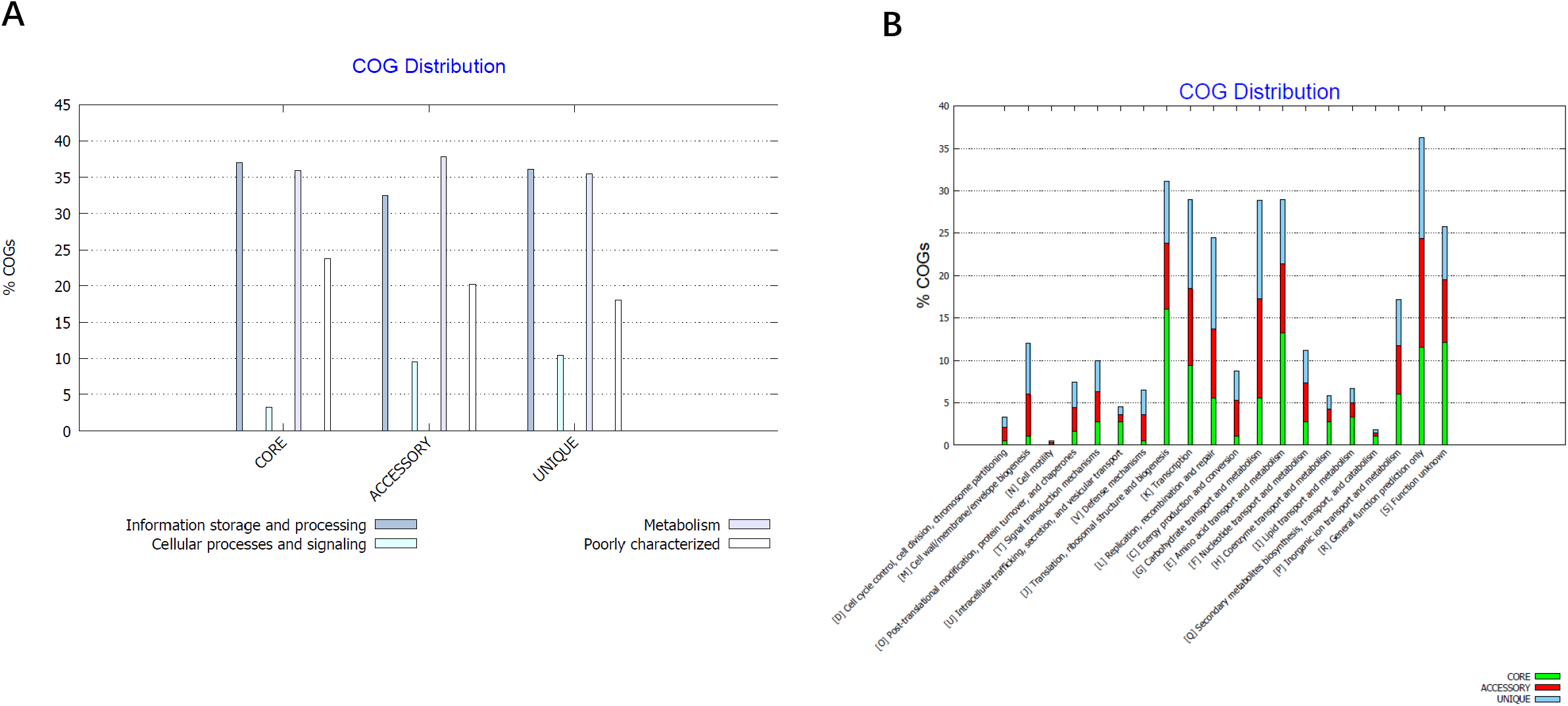
**A:** The general distribution of pan-genome annotated by using the COG database. Column plots show the distribution of pan-genome in general. **B:** The detailed distribution of pan-genome annotated by using the COG database. Stacked column plots show the distribution of pan-genome in detail.

**Supplementary Figure 7.**
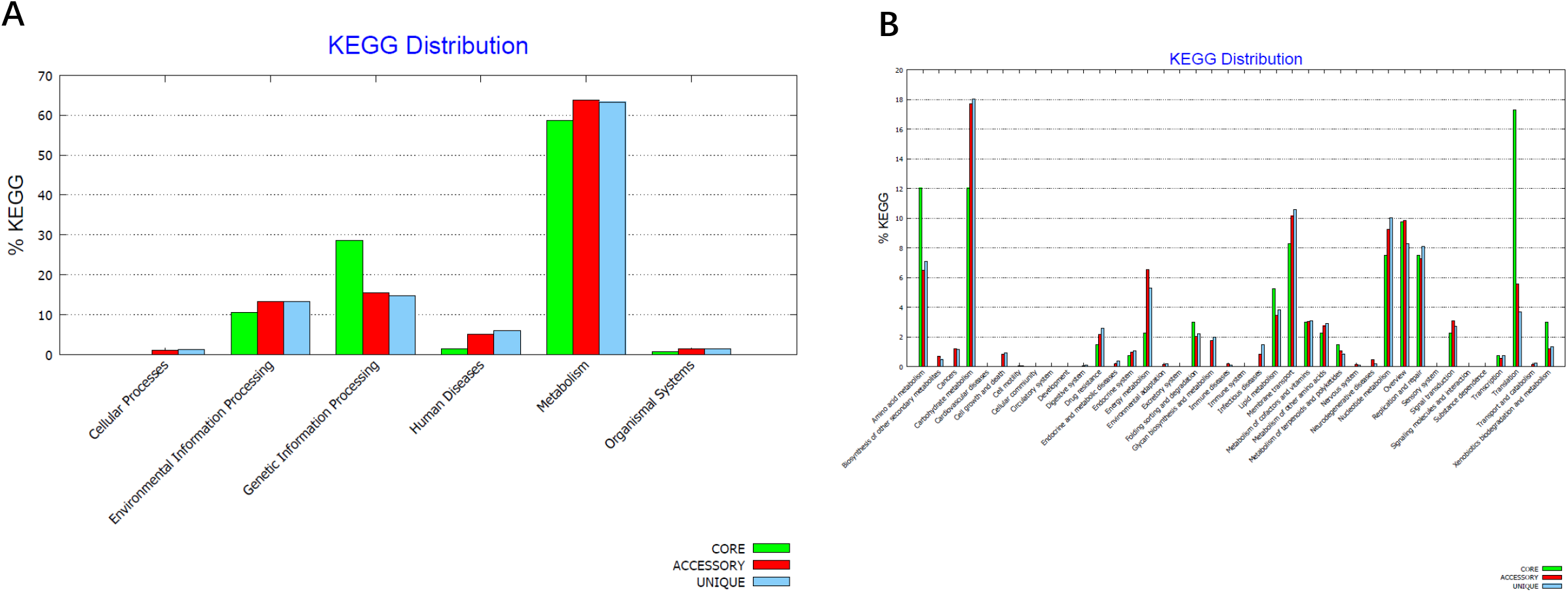
**A:** The general distribution of pan-genome annotated by using the KEGG database. Clustered column plots showing the distribution of pan-genome in general. **B:** The detailed distribution of pan-genome annotated by using the KEGG database. Clustered column plots showing the distribution of pan-genome in detail.

**Supplementary Fig 8.**
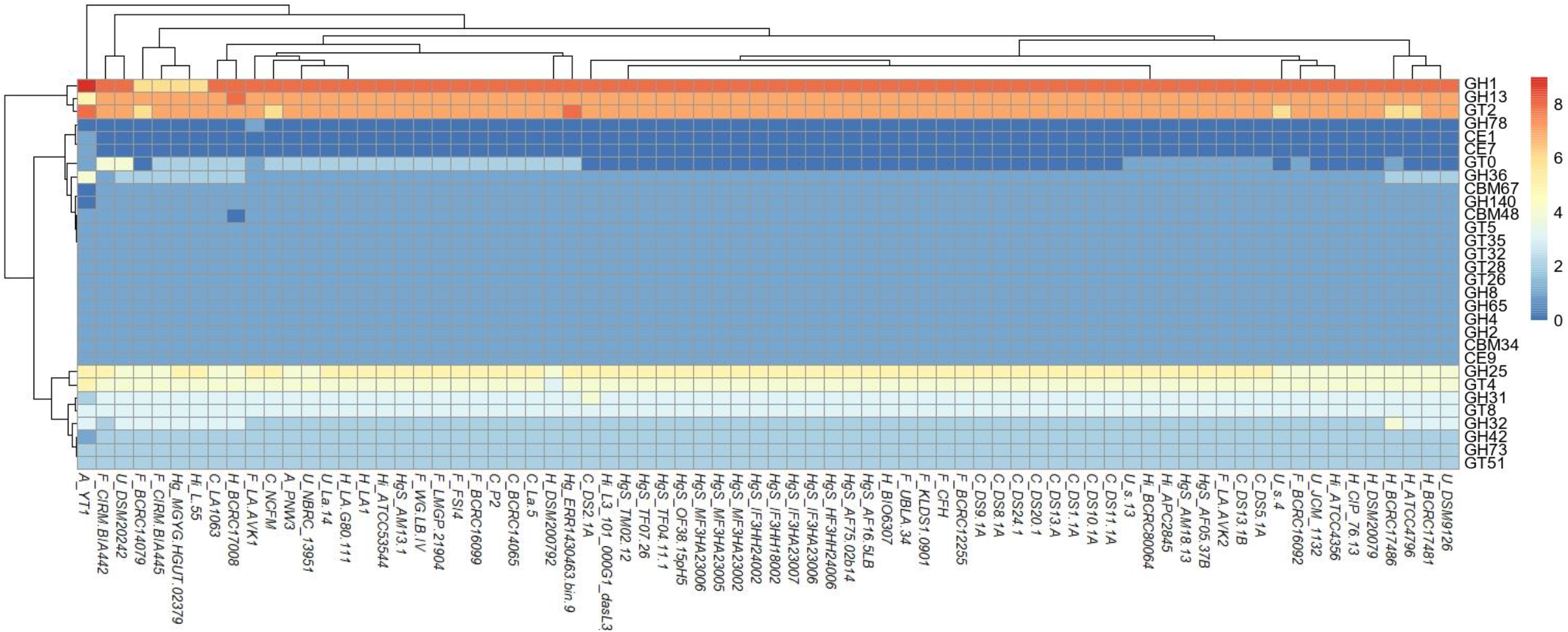
Carbohydrate utilization families identified in *Lactobacillus acidophilus*. Heatmap showing the distribution of carbohydrates utilization families.

