## Supplementary material for "Comparative Genome Analysis of *Lactobacillus acidophilus* Isolates from Different Ecological Niches": (Supplementary Table 5)The Genome Features of Lactobacillus.docx

| **Species** | **Sources** | **Average Genome Size (Mb)** | **Average GC%** | **Average CDS number** | **ANI range %** | **Pan genomes** | **Strict core genes** | **Reference** |
| --- | --- | --- | --- | --- | --- | --- | --- | --- |
| *Lactobacillus acidophilus* | Human gut, animal, food, commercial. | 1.97 | 34.61 | 1804 | 99.2 - 99.9 | 2329 | 1573 | This work |
| *Lactobacillus crispatus* | Human gut, vagi-l, oral cavity, animal. | 2.30 | - | 2245 | - | 3929 | 1224 | [1] |
| *Lactobacillus paracasei* | Human gut, food. | 2.97 | 46.40 | 2984 | - | 4200 | 1800 | [2] |
| *Lactobacillus mucosae* | Human, animal (piglets, dogs and cattles), food. | 2.11 | 47.96 | - | 95.3 – 99.9 | 8753 | 755 | [3] |
| *Lactobacillus ruminis* | Human, animal (bovines, dogs, porcine, horse), food. | 2.16 | 43.65 | 2309 | 96 - 99 | 1188 | 11188 | [4] |
| *Lactobacillus rhamnosus* | human gut, vagina, mouth, food. | 2.93 | 47.06 | 2838 | 97.2 – 99.9 | 190 | 8395 | [5] |
| *Lactobacillus gasseri* | Human gut. | 1.96 | 34.89 | 1936 | 93 - 99 | 1256 | 6535 | [6] |
| *Lactobacillus kefiranofaciens ZW3* | Kefir grains(food). | 2.11 | 37.7 | 2181 | - | - | - | [7] |
| *Lactobacillus helveticus* | Human gut, food. | 2.14 | 37.0 | 2151 | - | 998 | 3335 | [8] |

“-”: unknown
